# MCF-7 breast cancer cell line PacBio generated transcriptome has ~300 novel transcribed regions, un-annotated in both RefSeq and GENCODE, and absent in the liver, heart and brain transcriptomes

**DOI:** 10.1101/100974

**Authors:** Sandeep Chakraborty

## Abstract

Illuminating the ‘dark’ regions of the human genome remains an ongoing effort, a decade and a half after the human genome was sequenced - RefSeq and GENCODE being two of the major annotation databases. Pacific Biosciences (PacBio) has provided open access to the transcriptome of MCF-7, a breast cancer cell line that has provided significant therapeutic advancement in breast cancer research since the 1970s. PacBio sequencing generates much longer reads compared to second-generation sequencing technologies, with a trade-off of lower throughput, higher error rate and more cost per base. Here, this transcriptome was analyzed using the YeATS pipeline, with additionally introduced kmer based algorithms, reducing computational times to a few hours on a simple workstation. Out of ~300 transcripts that have no match in both RefSeq and GENCODE, ~250 are absent in the transcriptomes of the heart, liver and brain, also provided by PacBio. Also, ~200 transcripts are absent in a recent catalogue of un-annotated long non-coding RNAs from 6,503 samples (~43 Terabases of sequence data) [1], and among 2,556 novel transcripts reported in an experimental workflow RACE-Seq [2]. 65 transcripts have >100 amino acid open reading frames, and have the potential of being protein coding genes. ORF based annotation also identified few bacterial transcripts in the PacBio database mapped to the human genome, and one human transcript that has been annotated as bacterial in the NCBI database. The current work reiterates the under-utilization of transcriptomes for annotating genomes. It also provides new leads for investigating breast cancer by virtue of exclusively expressed transcripts not expressed in other tissues, which have the prospects of breast cancer biomarkers based on further investigations.

## Introduction

The initial estimates of 25,000 protein coding genes in humans has been moderated to about 19,000 recently [3]. This constitutes ~1% of the genome, but most of the 99% ‘dark’ genome plays significant regulatory roles in the cellular machinery [4]. The annotation of these regions is critical for correlating disease to genomic variants [5]. The two major independent annotation databases, periodically updated, are RefSeq [6] and GENCODE [7].

Pacific Biosciences (PacBio) sequencing [8] generates much longer reads compared to second-generation sequencing technologies [9], with a trade-off of lower throughput, higher error rate and more cost per base [10, 11]. The longer sequence lengths in PacBio compared to other sequencing methods might alleviate assembly issues associated with other methods with shorter read lengths [12, 13]. PacBio has provided open access to the transcriptome of the MCF-7 breast cancer cell line [14, 15]. There are currently two versions - one provided in 2013 and one in 2015. Two novel long non-coding RNAs were discovered in human mitochondrial DNA using the 2015 database by a different group [16].

The under-utilization of transcriptomes while annotating genomes [17–19] was recently emphasized for the walnut genome [20]. Here, the publicly available transcriptome of the MCF-7 breast cancer cell line (2013 version) was used to find novel human transcripts that are not annotated in current databases. ~300 transcripts are identified in the MCF-7 cells that have no annotation in the current RefSeq and GENCODE databases. Moreover, most of these transcripts are absent in heart, liver and brain transcriptomes also provided by PacBio. Also, ~200 transcripts are absent in a recent catalogue of un-annotated long non-coding RNAs (lncRNA) from 6,503 samples (~43 Terabases of sequence data) [1]. Furthermore, there no common transcripts with another experimental workflow RACE-Seq (rapid amplification of cDNA ends) and long-read RNA sequencing that reported 2,556 novel transcripts [2]. 65 of these transcripts have >100 long open reading frames (ORF), and might represent novel protein coding genes. A criteria comparing the amino acid frequency of the ORFs to the standard amino acid frequency found in human proteins is described in order to exclude proteins enriched for single amino acids which have repetitive sequences. Also, comparison of the ORFs to the BLAST ‘nr’ database highlights certain bacterial transcipts (Accid:WP_069187498.1) mapped to the human genome in the PacBio dataset, and one human protein (Accid:CPR56970.1) being annotated as a *Chlamydia trachomatis* protein in the NCBI database.

## Materials and methods

### GENCODE dataset

GENCODE release 25 was downloaded from https://www.gencodegenes.org/ (release date 07/2016). Two files - gencode.v25.transcripts.fa (n=200k) and gencode.v25.lncRNA transcripts.fa (n=27k) - were combined to create a single database(GENCODE. NTDB).

### RefSeq dataset

The RefSeq database was created from https://www.ncbi.nlm.nih.gov/nuccore choosing mRNA, rRNA, cRNA, tRNA and ncRNA sequences (FILE:mrna.refseq.160k.fa, n=161k, REFSEQ.NTDB).

### PacBio dataset

The MCF-7 transcriptome was obtained from http://www.pacb.com/blog/data-release-human-mcf-7-transcriptome (2013 version). There is another updated version made available in 2015, that has not been analyzed in this study. The PacBio dataset for human heart, liver and brain transcriptomes is available at http://datasets.pacb.com.s3.amazonaws.com/2014/Iso-seq_Human_Tissues/list.html and provides ‘a dataset containing the full-length whole transcriptome from three diverse human tissues (brain, heart, and liver). The updated version of the Iso-Seq method incorporates the use of a new PCR polymerase that improves the representation of larger transcripts, enabling sequencing of cDNAs of nearly 10 kb in length. The inclusion of multiple sample types makes this dataset ideal for exploring differential alternative splicing events’ [21]. The transcripts from all three tissues were concatenated to create a single file (FILE:PacBioHLB.fa, n=23309). The transcripts have been renamed to allow Unix style filenames.

### kmer analysis

The MCF-7 transcriptome was 30kmer compared to the GENCODE.NTDB and REFSEQ.NTDB. This step removes all transcripts that have at least a 30 long sequence (ignoring repetitive sequences) common in the respective annotation databases. This is a conservative step that just reduces the search database, since a transcript ignored as mentioned above might still not match exactly to any entry in the annotation database. The ‘unmatched’ transcipts are now BLAST’ed [22] to GENCODE.NTDB and REFSEQ.NTDB to have a complete alignment. This results in two sets of transcripts. The first set has no matches in the database, and are definitely not annotated. The second set has homologous entries in the database, but need to be manually checked for homology. It is seen that entries which are <96% homologous are mostly novel annotations. Open reading frames were obtained ‘getorf’ program from the EMBOSS suite [23]. Computational runtimes are very modest, and takes a few hours on a personal workstation.

## Results and discussion

### No match in the RefSeq and GENCODE databases

First, transcripts that shared a 30kmer sequence with any annotated sequence in the GENCODE and RefSeq databases were removed. Next, a BLAST comparison of transcripts that do not share a 30kmer sequence in the respective GENCODE and RefSeq databases provided transcripts which have no matches in the GENCODE.NTDB (FILE:notinGENCODE.list, n=346) and REFSEQ.NTDB (FILE:notinREFSEQ.list, n=418). This corroborates a recent study that found GENCODE to have better coverage [24]. There are 252 transcipts common to these sets.

### Partial homology in the RefSeq and GENCODE databases

The BLAST comparison also provided a list of transcripts with partial homology to the RefSeq and GENCODE databases. Transcripts with >96% homology are probably annotated, and excluded. There are 42 transcripts that have <96% homology with respect to both databases, and thus can be considered as not being annotated (FILE:notInBoth.96homology.list). These transcripts require manual inspection.

### Comparison to the PacBio transcriptome of the heart, liver and brain

Next, these transcripts were combined (FILE:notInBoth.list, n=294, Fig 1) and compared to the PacBio provided dataset containing the full-length whole transcriptome from three diverse human tissues (brain, heart, and liver, FILE:PacBioHLB.fa). There were 264 transcripts that were not present in these tissues (FILE:notin.HLBdatabase.list), thus representing transcripts exclusive to the MCF-7 cell line (at least with respect to the brain, heart, and liver).

**Figure 1:**
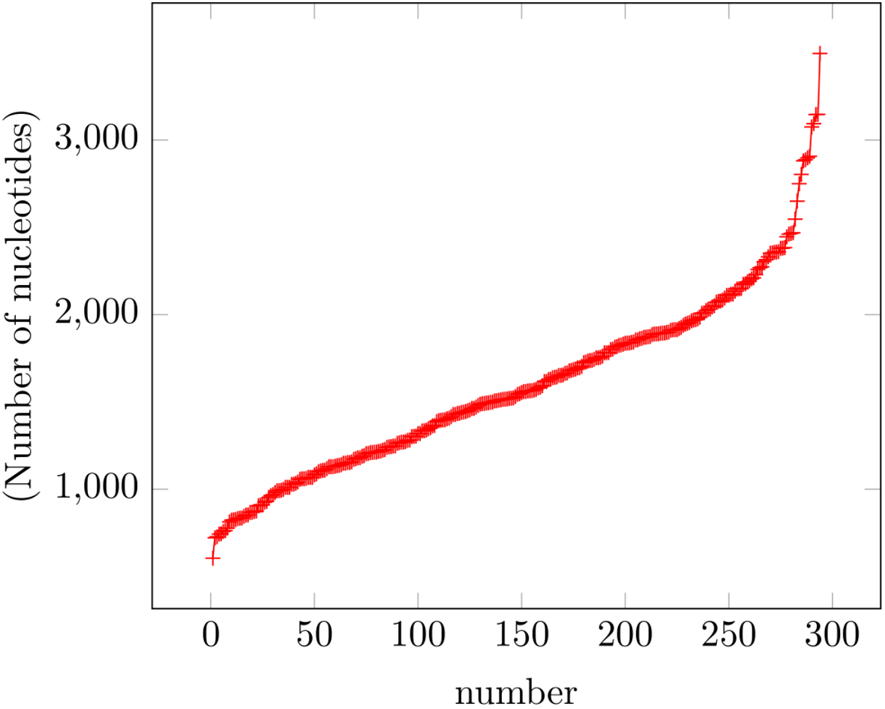
Length of the 294 transcripts not found in RefSeq and GENCODE databases: Shortest is ~600 nt long, while the longest is ~3500 nt. In contrast the mitranscriptome.org database has ~12k sequences ranging from 300 to 985680 nt.

### Comparison to the mitranscriptome.org database

The relevance of lncRNAs in cancer studies has long been established [25]. A recent work classified 58,648 genes as lncRNAs after a comprehensive analysis of ‘7,256 RNA-Seq libraries from tumors, normal tissues, and cell lines comprising over 43 terabases of sequence from 25 independent studies’ [1]. Supplementary Table 10 (http://www.nature.com/ng/journal/v47/n3/full/ng.3192.html#supplementary-information) provides the genomic coordinates of these lncRNAs, which was extracted using ‘http://genome.ucsc.edu/cgi-bin/das/hg19/dna? segment=CHR:start, end’. This information is also provided in http://mitranscriptome.org/. Follow-up work on this database demonstrated the importance of specific lncRNAs to breast cancer [26]. There are ~12k sequences ranging from 300 to 985680 nucleotides. Comparison of the 294 transcripts (FILE:notInBoth.list) to these sequences identified common transcripts (FILE:mitranscriptome.same, n=52) and unannotated transcripts (FILE:mitranscriptome.NOTTHERE, n=242). Unannotated transcripts include those that have homology (> BLAST bitscore 100) but map to a different chromosome, as well as those that have no homologs.

### Comparison to the RACE-seq novel transcripts

Another experimental workflow RACE-Seq (rapid amplification of cDNA ends) and long-read RNA sequencing reported 2,556 novel transcripts [2]. The genomic coordinates are obtained from https://public_docs.crg.es/rguigo/Papers/uszczynska/RACE-Seq/ (FILE:phase6-clean.bed, n=2486). None of the novel transcripts identified here were reported in this study (FILE:RACESEQ.THERE).

### Identification of possible protein coding genes

Transcripts encoding open reading frames (ORF) longer than 100 amino acids are not typically considered as lncRNAs [27]. There are 65 transcripts with >100 long ORFs (FILE:notInBoth.BRCA.list.ORF.100.fa). The amino acid frequency (AAF) of these ORFs can be compared to the combined AAFs found in all human proteins (Fig 2) to select more likely protein coding genes.

**Figure 2:**
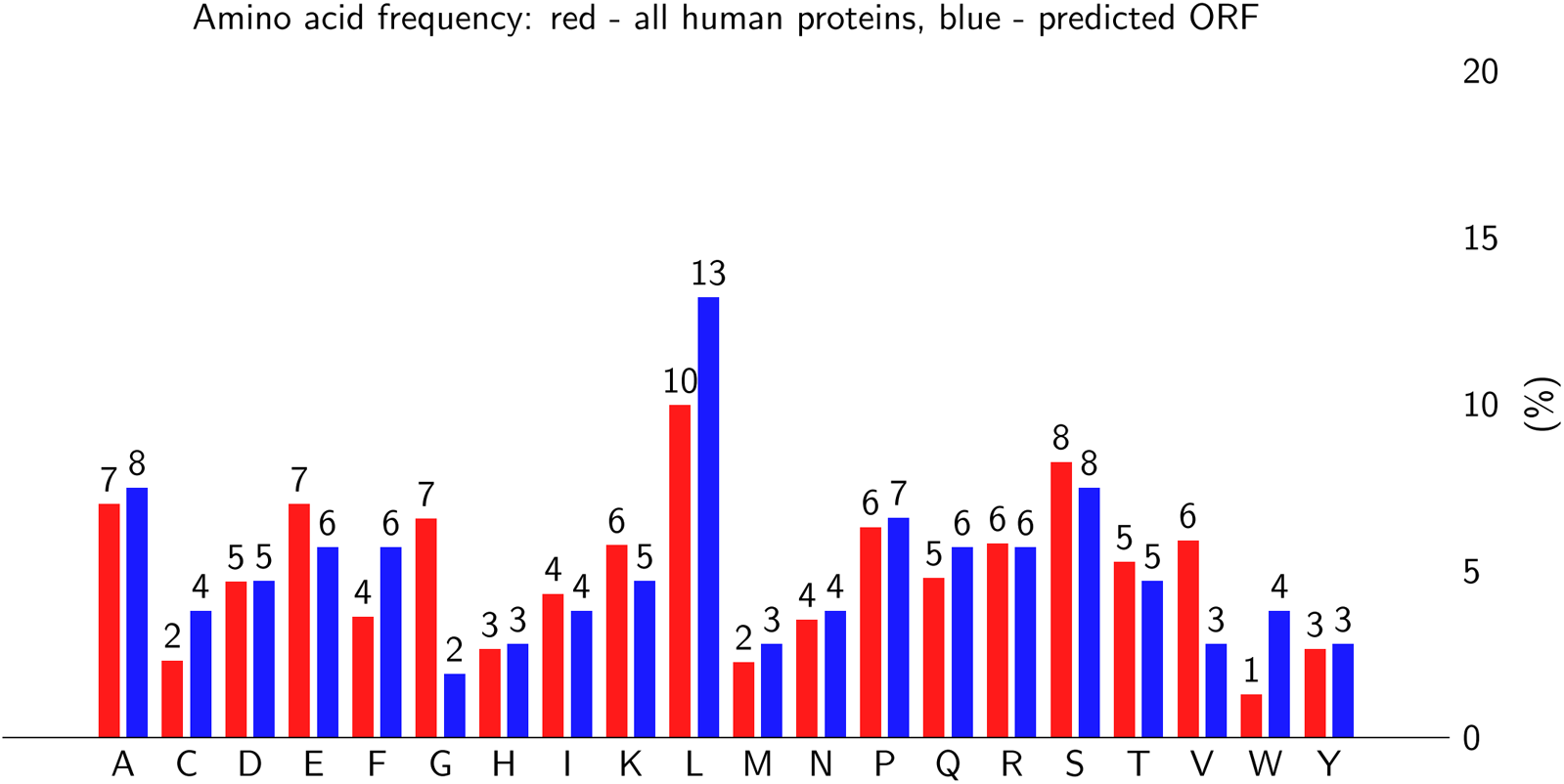
Amino acid frequency (AAF) in human proteins: The cumulative AAF (in red) is obtained from the ncbi RefSeq human proteins (n=80k). The AAF for the ORF from the transcript CHR5.67101977_67106707+.I3_C6784.F3P34.3147.RT_6 (length=106 aa) is in blue.

### Misannotated transcripts in the PacBio dataset and NCBI dataset

These ORFs were BLAST’ed to the complete ‘nr’ database (Table 1). This highlighted that bacterial transcipts from *Streptococcus pneumoniae* and *Mycobacterium tuberculosis* have being annotated as human transcipts in the PacBio database. For example, CHR16.46389879_46398888._I3_C30162.F2P21.3075.RT has a 89% homology to chr16 (which explains the annotation), and no homology to *S. pneumoniae*. However, it encodes an ORF (132 aa long) which has a 100% identity to a protein from *S. pneumoniae* (Accid:WP_069187498.1). This emphasizes the importance of using ORF based annotation for transcripts with low homology [28]. Another human protein (Accid:CPR56970.1) has been mistakenly annotated as a *Chlamydia trachomatis* protein in the NCBI database, based on a vaginal swab transcript (Accid:CSTN01000036.1, 2920 nt) that has a 99% homology to the human chromosome 1.

**Table 1:** Transcripts with ORFs >100 aa long that match to the ‘nr’ database: Transcript 1, 2 and 4 are bacterial transcripts, demonstrating the necessity of removing metagenomic transcripts prior to analysis. Transcript 8 is misannotated as protein from *Chlamydia trachomatis*. The transcript (Accid:CSTN01000036.1) linked to the protein (Accid:CPR56970.1) matches to the human genome, and not to *Chlamydia trachomatis*. A cutoff of BLAST bit score (BBS) of 100 was used. The other ~ 90 transcripts might represent novel protein coding genes or lncRNAs.

| Idx | Transcript | Accid | Description | BBS |
| --- | --- | --- | --- | --- |
| 1 | CHR16.46389879_46398888_I3_C30162.F2P21.3075.RT_29 | WP_069187498.1 | [*S. pneumoniae*] | 247 |
| 2 | CHR16.46386138_46388586_I2A_C135052.F1P32.2469.RT_26 | CJE27995.1 | [*S. pneumoniae*] | 216 |
| 3 | CHR19.39155468_39158343+.I2A_C37015.F3P20.2881.RT_18 | EAW56812.1 | [*H. sapiens*] | 197 |
| 4 | CHR16.46386138_46388586_I2A_C135052.F1P32.2469.RT_36 | CKT10751.1 | [*M. tuberculosis*] | 178 |
| 5 | CHR15.99344096_99385537+.I1D_C21966.F4P10.1565.RT_15 | EAX02227.1 | [*H. sapiens*] | 140 |
| 6 | CHR11.70996950_70998864+.I1C_C72675.F1P38.1921.RT_32 | AIE45854.1 | [synthetic construct] | 119 |
| 7 | CHR2.12841430_12856867_I1D_C30377.F5P15.1894.RT_31 | XP_011747760.1 | [*M. nemestrina*] | 112 |
| 8 | CHR1.145370688_145372138+.I1A_C27600.F1P13.1454.RT_5 | CPR56970.1 | [*C. trachomatis*] | 105 |
| 9 | CHR7.50518150_50519652+.I1D_C31040.F3P6.1510.RT_2 | EAW60974.1 | [*H. sapiens*] | 101 |

## Conclusions

The current work highlights the ‘low hanging fruits’ still available for widely researched diseases like breast cancer, at very low computational costs. The technological advancement provided by PacBio sequencing might be responsible for identification of these transcripts, that has eluded detection thus far. It also highlights the necessity of metagenomic filtration prior to analysis.

## Revisions from version 1

1. The number of transcripts having ORFs >100 aa is updated to 65 from ~100. The program for analyzing was considering mutliple ORFs from the same transcript previously - this has been fixed.
2. It was previously mentioned that there were two transcripts that were common in the ~2500 novel transcripts identied in the RACE-seq study. The homology of these matches is too low to classify these as matches.

